# Glue genes are subjected to diverse selective forces during Drosophila development

**DOI:** 10.1101/2021.08.09.455518

**Authors:** Flora Borne, Rob J. Kulathinal, Virginie Courtier-Orgogozo

## Abstract

Molecular evolutionary studies usually focus on genes with clear roles in adult fitness or on developmental genes expressed at multiple time points during the life of the organism. Here, we examine the evolutionary dynamics of Drosophila glue genes, a set of eight genes tasked with a singular primary function during a specific developmental stage: the production of glue that allows animal pupa to attach to a substrate for several days during metamorphosis. Using phenotypic assays and available data from transcriptomics, PacBio genomes, and genetic variation from global populations, we explore the selective forces acting on the glue genes within the cosmopolitan *D. melanogaster* species and its five closely related species, *D. simulans, D. sechellia, D. mauritiana, D. yakuba*, and *D. teissieri*. We observe a three-fold difference in glue adhesion between the least and the most adhesive *D. melanogaster* strain, indicating a strong genetic component to phenotypic variation. These eight glue genes are among the most highly expressed genes in salivary glands yet they display no notable codon bias. New copies of *Sgs3* and *Sgs7* are found in *D. yakuba* and *D. teissieri* with the *Sgs3* coding sequence evolving rapidly after duplication in the *D. yakuba* branch. Multiple sites along the various glue genes appear to be constrained. Our population genetics analysis in *D. melanogaster* suggests signs of local adaptive evolution for *Sgs3, Sgs5* and *Sgs5bis* and traces of selective sweeps for *Sgs1, Sgs3, Sgs7* and *Sgs8*. Our work shows that stage-specific genes can be subjected to various dynamic evolutionary forces. (249 words)

**Significance statement:** Drosophila larvae produce a glue to stick themselves to a substrate for several days during metamorphosis. Here we observe wide variation in stickiness among *Drosophila melanogaster* strains and we analyze the molecular evolution of eight glue genes. We find several recent gene duplications and heterogenous rates of evolution among these genes.

## Introduction

Our understanding of evolutionary patterns and processes in multicellular eukaryotes derives primarily from observations and analyses of adult stages. For example, in Gephebase, a database that compiles genes that have been found to contribute to evolutionary changes in animals and plants (Courtier-Orgogozo et al., 2020), about 95% of the phenotypic traits refers to the adult stage while only 5% corresponds to earlier developmental stages. Yet a large component of an individual’s relative fitness may occur before the adult stage. Here we use a simple model system to study the influence of evolutionary forces on genes whose function appears to be restricted to a specific developmental stage, the genes encoding for the glue that attaches the animal to external substrates during the pupal stage in Drosophila. In Diptera, development transitions through several larval stages followed by a pupal stage during which metamorphosis occurs. During the pupal stage, insects are particularly vulnerable because they are mostly immobile. In Drosophila, pupation site choice during the larval stage depends on various environmental conditions such as light (Rizki and Davis Jr, 1953), temperature (Schnebel and Grossfield, 1992), and humidity (Sokolowski, 1980) but is also influenced by intra- and interspecific competition (Beltramí et al., 2012; Da Silva et al., 2019). Insects have evolved different mechanisms to protect pupa from predation such as coloration, production of toxins, aggregation (Lindstedt et al., 2019), and pupal adhesion (Borne et al., 2021). For example, in both *Drosophila simulans* and *D. suzukii*, attached pupae survive longer in laboratory assays in the presence of ants compared to manually detached pupae (Borne et al., 2021). The ability of pupae to successfully adhere to a diverse array of surfaces thus represents an important trait subjected to selection in fruit flies.

In *D. melanogaster*, a glue is produced by late third instar larvae from their salivary glands a few hours before pupariation (Duan et al., 2020). The glue polymerizes quickly after expectoration and allows the animal to attach firmly to various substrates (Fraenkel and Brookes, 1953). The glue is primarily composed of eight proteins encoded by the genes *Sgs1, Sgs3, Sgs4, Sgs5, Sgs5bis, Sgs7, Sgs8*, and *Eig71Ee* (Korge, 1975, 1977; Da Lage et al., 2019). The glue genes are located within puffs of polytene chromosomes and have been a premier model for studying the mechanisms involved in the activation of gene expression by ecdysone in the 70’s (Akam et al., 1978; Korge, 1977, 1975). Glue proteins can be classified into two groups. One group comprises Sgs1, Sgs3, Sgs4, and Eig71Ee, which are mainly composed of repeat sequences rich in proline, serine and threonine, and are highly O-glycosylated, suggesting that they may interact with water to rehydrate the glue during the expectoration process to lubricate the lumen of the glands and the mouth (Farkaš, 2016). They may also contribute to glue adhesion by interacting with the substrate (Farkaš, 2016). The glue has been shown to adhere relatively strongly to polarizable substrates that may interact with the negative charges of sugar components and the positive charges of amino acid components of the glue (Borne et al., 2020). Furthermore, Sgs1, Sgs3, and Eig71Ee belong to the mucin family as they are characterized by poorly conserved extended regions of repeated sequences containing prolines and glycosylated serines or threonines (Syed et al., 2008, Da Lage et al., 2019;). As with other mucins, these proteins could harbor antimicrobial functions (Bakshani et al., 2018; Syed et al., 2008). The other group of glue proteins comprises Sgs5, Sgs5bis, Sgs7, and Sgs8, which are shorter and more ordered proteins that may interact with the other disordered glue proteins to prevent protein aggregation and allow the secretion of the glue (Farkaš, 2016). They could also have a role in adhesion. So far, the functions of the different glue proteins have not been assessed.

In a previous study, glue genes were identified in other Drosophila species via sequence similarity with annotated *D. melanogaster* genes and, so far, they have only been found within the Drosophila genus, likely due to rapid divergence of genetic sequences (Da Lage et al., 2019). In Drosophila, glue genes have undergone multiple gains and losses of copies as well as extensive genetic changes in their coding sequences (Da Lage et al., 2019; Farkaš, 2016). In particular, repeat regions of *Sgs1, Sgs3, Sgs4*, and *Eig71Ee* have evolved rapidly in terms of the number of repeats as well as the motif sequenceof these repeats (Da Lage et al., 2019). A first attempt was made to study the evolutionary rate of the glue genes but was primarily limited to the glue genes without repeats due to the limited quality of the available genome assemblies at the time (Da Lage et al., 2019).

Recent advances in long read sequencing (PacBio or Oxford Nanopore) now make it possible to utilize higher quality genomes with reliable sequences spanning across multiple repeats. High quality assemblies have been recently generated in *D. melanogaster* and closely related species (e.g., Chakraborty et al., 2019; Kim et al., 2021) and can be applied for the study of repeat-laden genes such as glue genes. Glue genes seem to be expressed exclusively in the salivary glands over a relatively short period of time during the larval stage (Andres et al., 1993; Duan et al., 2020; Li and White, 2003), with the exception of Eig71Ee which is also expressed in hemocytes and in the gut where it is probably involved in immunity and clotting (Korayem et al., 2004). Their function thus appears to be limited to glue properties where they may play an important and very specific role in the fly’s ultimate survival. This set of tissue- and developmental stage-specific genes together with our recently developed phenotypic assay to quantify pupal adhesion (Borne et al., 2020) provides a promising model to understand how patterns of genetic variation are related to phenotypic variation in adhesion as well as the role of adaptation in early metamorphic stages.

In this study, we investigate the phenotypic variation in pupal adhesion and the genetic variation of glue genes in Drosophila. We apply a force assay (Borne et al. 2020) on individual pupae from a set of 12 inbred *D. melanogaster* lines originating from different geographic locations (Chakraborty et al., 2019) as well as three sister species to survey differences in pupal adhesion. Using fifteen high-quality PacBio genomes that correspond to these phenotyped lines and three additional lines from *D. melanogaster*, as well as high-quality PacBio genomes of five sister species, we then investigated the evolutionary dynamics of these glue genes. We observed low levels of codon bias on these highly expressed genes, discovered several gene duplications, and identified rapidly evolving lineages. Putative sites with signals of negative and positive selection were found in these glue genes among *D. melanogaster* lines as well as global populations from the Drosophila Genome Nexus (Lack et al., 2015). Such stage- and tissue-specific genes provide an excellent model to further study the evolutionary dynamics underlying fitness effects during a particular stage of an organism’s life cycle.

## Results

### Pupal adhesion varies up to three-fold between *D. melanogaster* lines

We compared pupal adhesion of twelve *D. melanogaster* lines from geographically diverse regions, isogenized in the laboratory and maintained over decades, by measuring the force at which pupae were detached from glass slides (Figure 1). High variance is observed among individuals from the same strain but still 26 % of the total phenotypic variance is explained by strain (ANOVA F = 11.96, df = 15, p<2e-16). Median adhesion force varies by three-fold across lines, from 124.45 mN (SD=69 mN) in the A5 strain to 377.32 mN (SD=131 mN) in A7. In particular, we could distinguish a cluster of low adhesive lines composed of A5 and B4 strains and a cluster of highly adhesive lines composed of A7 and B6 (ANOVA followed by multiple pairwise comparisons, p<0.05, Figure 1). We recorded room temperature and humidity during the adhesion assays to examine potential effects of environmental factors. In our experiments, humidity negatively correlates with temperature (Pearson correlation, t=-16.05, df=382, cor=- 0.63, p<2.2e-16), humidity has no effect on adhesion (univariate linear model, F=0.14, df=382, R^2^=0.0004, p=0.7) and temperature may have a slight negative effect on adhesion (univariate linear model, F=5.44, df=382, R^2^=0.01, p=0.02). To test whether differences in adhesion between lines were due to a difference in the surface of contact between the glue and the substrate, areas of the prints left by the pupae on glass slides after detachment were measured. We found that contact areas vary significantly between lines (ANOVA F=14.05, df=11, p<2e-16; Figure S1) but no correlation was found between the force and the pupa-substrate contact area (sma regression, p=0.30), which suggests that differences in adhesion between genotypes are not due to changes in the amount of glue produced by larvae but rather in glue adhesive properties. It could also suggest that glue prints left on glass slides after pupa removal may not be a good proxy for estimating the surface of contact between the glue and the substrate. We quantified levels of pupal adhesion in three other species from the *D. melanogaster* complex and found no difference between species, except for *D. mauritiana* which has a lower adhesion (ANOVA followed by pairwise comparisons: *D. simulans* - *D. mauritiana*: p<0.05; *D. melanogaster* - *D. mauritiana:* p<0.01; Figure 1).

**Figure 1.**
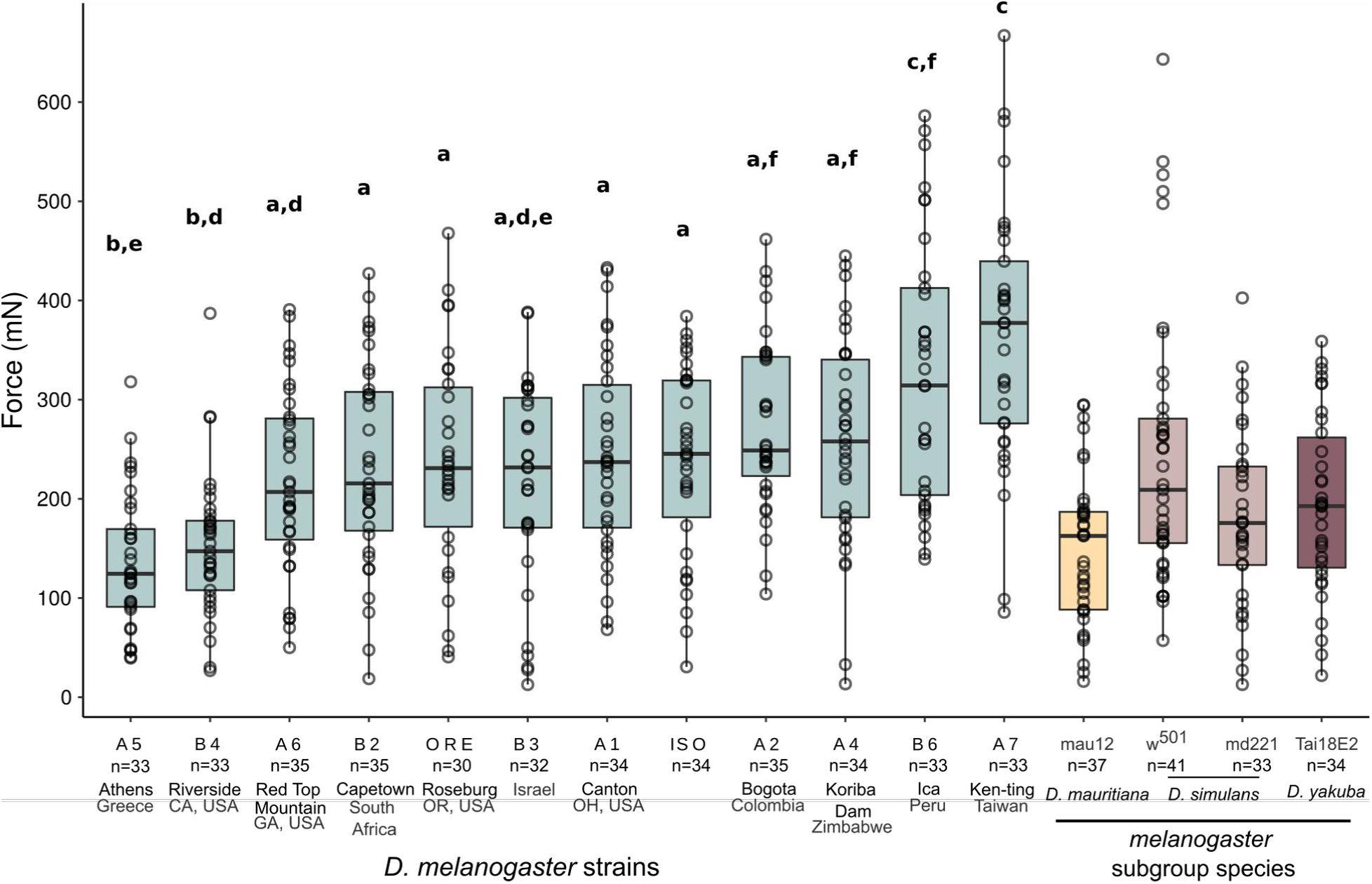
Variation of pupal adhesion force across global *Drosophila melanogaster* lines and closely related species of the *D. melanogaster* subgroup. The force required to detach a pupa naturally attached to a glass slide was measured on individual pupae. n indicates the total number of pupae measured for each strain. Boxes define the first and third quartiles and the black horizontal line represents the median. The upper vertical line extends to the largest value no further than 1.5 * IQR from the box and the lower vertical line extends to the smallest value at most 1.5 * IQR. (IQR: inter-quartile range is the distance between the first and the third quartiles). Boxes are shaded by color to differentiate each species. An ANOVA followed by all pairwise comparisons after Tukey correction (p < 0.05) was performed on the set of *D. melanogaster* strains. Lines that are not significantly different from each other share a letter.

### Glue genes are the most highly expressed genes in salivary glands but they display no notable codon bias

We used previously published transcriptome data from *D. melanogaster* OregonR salivary glands dissected at the wandering third instar larval stage (Graveley et al., 2011) to examine the level of expression of glue genes. We found that our eight glue genes are among the ten most highly expressed genes, with expression levels being about 1,000 to 300,000-fold greater than the median (Figure 2A, see Dataset1.ods in Dryad). We then measured codon adaptation index (CAI) for *D. melanogaster* genes expressed in the salivary glands of wandering larvae. CAI measures the deviation of codon usage of a given gene compared to a reference set of genes and is expected to correlate with gene expression levels (Sharp and Li, 1987). Whereas a positive correlation between gene expression and CAI can indeed be identified with the full set of genes expressed in wandering larvae salivary glands (Kendall correlation, z=4.62, τ=0.10, p=3.85e-06), the glue genes do not appear to be part of this global trend and are generally less biased than their co-expressed genes, even though they are among the most expressed genes (Figure 2A). When excluding the repetitive regions, we found CAI (*Sgs1*:0.648, *Sgs3*: 0.659, *Sgs4*: 0.676, *Eig71Ee*: 0.649) to be comparable to CAI estimated on the full coding sequence. Overall, we distinguish two types of highly expressed genes in the salivary glands of L3 wandering larvae: a first group with high CAI (encoding for ribosomal proteins, cytoskeleton components, etc.) and a second group with low to medium CAI, i.e., little codon bias (glue genes and genes of unknown function) (Table S1). Among codons that code for threonine and proline, the two most abundant amino acids found in repeat regions, U- and A- ending codons were mainly used in *Sgs1, Eig71Ee*, and *Sgs4* whereas codons ending in C were primarily used in *Sgs3* (Figure 2B). This difference could be explained either by different constraints on the nucleotide repeats of *Sgs3* compared to *Sgs1* and *Eig71Ee* or by concerted evolution of the repetitive sequences within each gene via unequal crossover, expansion or contraction of the array, or gene conversion (Elder Jr and Turner, 1995).

**Figure 2.**
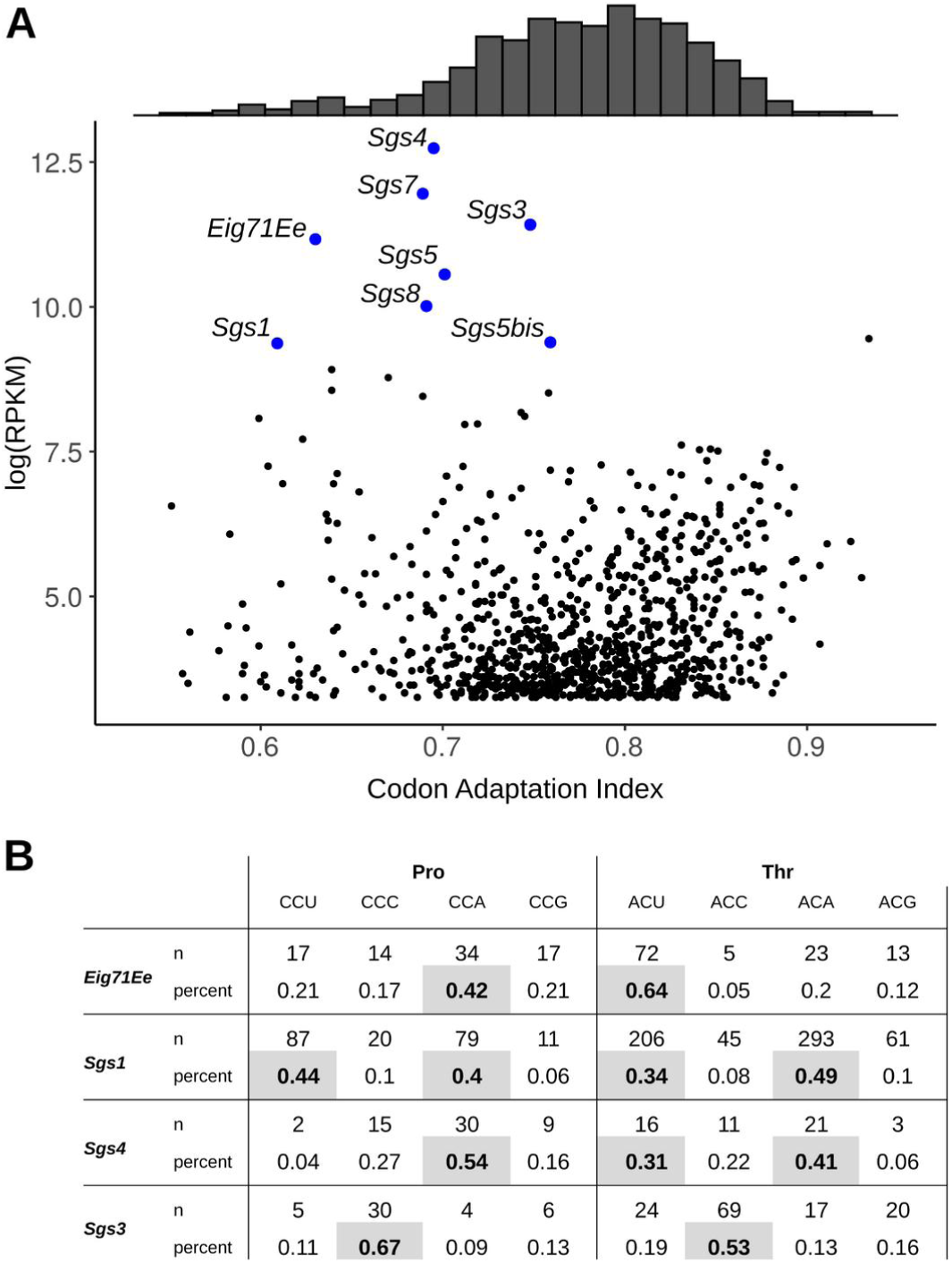
Codon usage of the genes expressed in the salivary gland of third instar larvae in *D. melanogaster*, highlighting the glue genes. A) Relationship between expression level (log(RPKM)) and codon adaptation index (CAI) of *D. melanogaster* genes expressed in the salivary glands of wandering larvae. CAI histogram is represented above. B) Codon usage of *Sgs1, Sgs3, Sgs4* and *Eig71Ee* for the two most abundant amino acids in repeat regions, proline and threonine. Codons with fractions greater than 0.3 are highlighted in grey.

### *Sgs3* and *Sgs7* duplicated recently and *Sgs3* coding sequence evolved rapidly after duplication in the *D. yakuba* branch

Using well-assembled PacBio genomes and synteny, we identified and annotated glue genes in *D. melanogaster* (A1 strain) and five closely related species (*D. simulans, D. mauritiana, D. sechellia, D. yakuba* and *D. teissieri*) (Figure 3A). Interestingly, the *Sgs5* gene is missing in *D. mauritiana*. We identified two *Sgs3* copies in *D. yakuba*, three *Sgs3* copies in *D. teissieri*, and two *Sgs7* copies in both species. The extra copies of *Sgs3* were not found in a previous study using genome assemblies that derived from Illumina short reads (Da Lage et al., 2019). The *Sgs3bis* genes are located at exactly the same region of the genome in *D. yakuba* and *D. teissieri* indicating that they originate from the same duplication event (Figure S2A). The same observation was found for their *Sgs7bis* genes. At the nucleotide level, the *Sgs7bis* and *Sgs7* genes from *D. teissieri* are 99.1% identical (2 SNPs difference) and the *Sgs7bis* and *Sgs7* genes from *D. yakuba* are 100% identical, but the *Sgs7/Sgs7bis* genes are 13% divergent between *D. yakuba* and *D. teissieri* (Figure S2B). This indicates that *Sgs7bis* and *Sgs7* derive from a unique duplication event preceding *D. yakuba* - *D. teissieri* divergence which was followed by gene conversion in both species.

**Figure 3.**
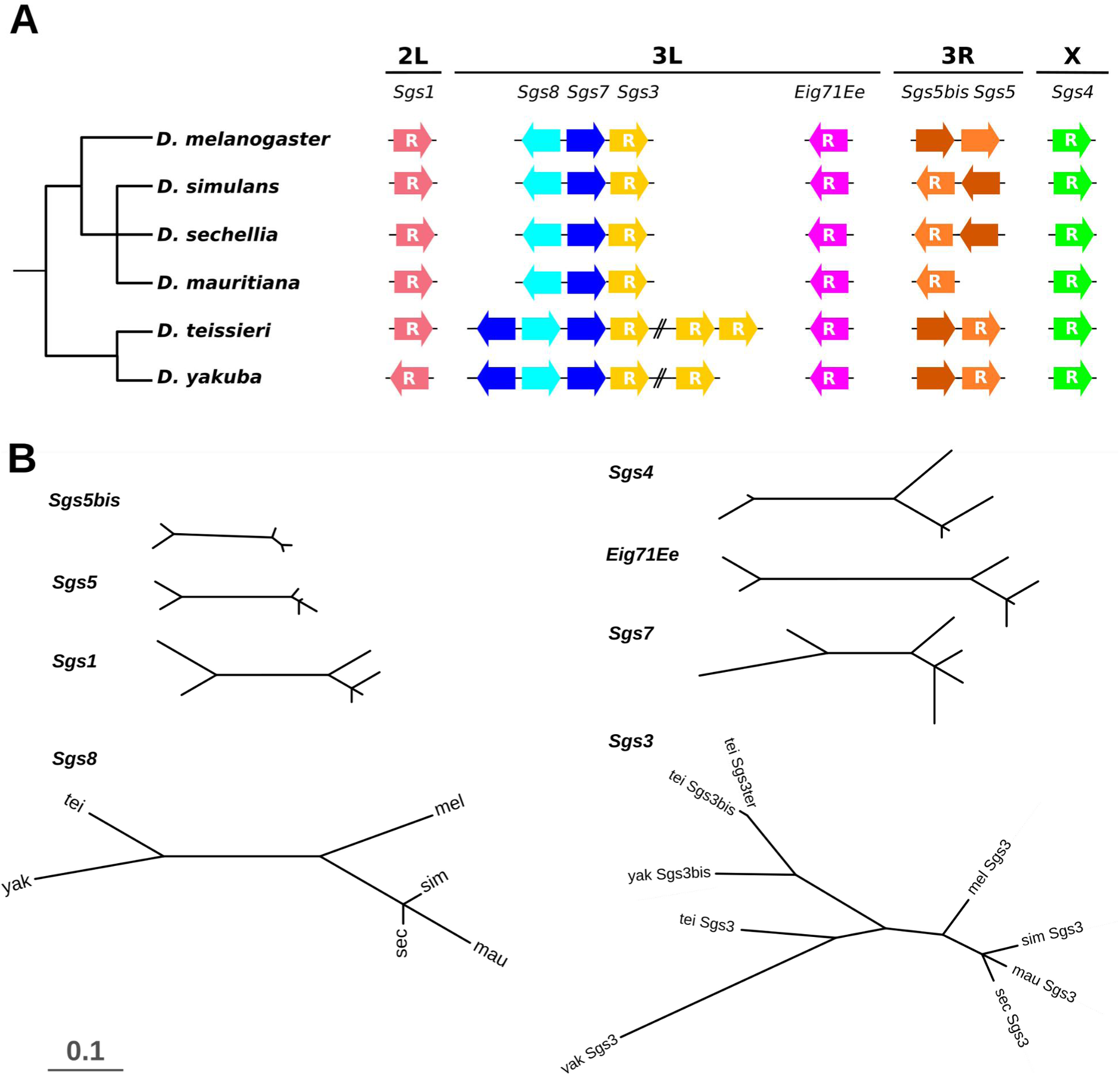
Evolution of Drosophila glue genes across species. A) Evolution of the number and orientation of glue genes. Species tree is shown on the left and the organization of the glue gene clusters on the right as in Da Lage et al. (2019). Corresponding strains from top to bottom species: *D. melanogaster*: A1; *D. simulans*: w^501^; *D. sechellia*: sec25; *D. mauritiana*: mau12; *D. teissieri*: GT53w; *D. yakuba*: NY73PB. Gene sizes and distances are not to scale. “R” means that internal repeats are present. B) Nonsynonymous substitution rate (dN) trees. Branch lengths are proportional to nonsynonymous substitution rates calculated by PAML. Species are positioned similarly in all trees as shown in the *Sgs8* dN tree: *D. yakuba* (yak), *D. teissieri* (tei), *D. melanogaster* (mel), *D. simulans* (sim), *D. sechellia* (sec), D. mauritiana (mau). The scale bar indicates the nonsynonymous substitution rate (0.1 substitution/site).

To test for lineage-specific selection, *Sgs3* in which the *D. yakuba* branch evolved more rapidly than other branches (Figure 3B, model statistics in Table S2), we used PAML for each *Sgs* gene (including *Sgs3, Sgs3bis* and *Sgs3ter* for *Sgs3* and a single *Sgs7* gene as *Sgs7* and *Sgs7bis* are basically identical sequences) and compared rates of evolution of the glue genes by performing nonsynonymous substitution rate trees. Since many repeats were too divergent, regions containing repeats were excluded. We found evidence of lineage-specific selection for The same trend was observed in *Sgs7* (Figure 3B). This rapid evolution may suggest adaptation of the glue in this branch. *Sgs8* seems to evolve more rapidly in all branches (dN tree length=0.50) than the other glue genes (dN tree length between 0.12 and 0.39 except for *Sgs3*: dN=0.59, Figure 3B), indicative of less functional constraints on its protein.

### Glue genes display signs of purifying selection

We used the Fixed Effects Likelihood (FEL) framework from HyPhy to test for sites under positive and negative selection across the six Drosophila species (Table 1). Again, regions containing repeats were excluded because they were too divergent to align between species. As the species set used is relatively small (n=6), the analysis may lack the power to detect weak signals of selection. We found multiple sites under negative selection in each glue gene (Table 1, Figure S3, see Dataset1.ods in Dryad). These sites were not clustered in specific regions of the protein-coding landscape but rather spread along the sequences, in both the peptide signal region and the regions located 5’ and 3’ of the repeat regions. This suggests that functional constraints exist throughout the extent of each glue protein. We found one site under positive selection in *Sgs4* which corresponds to a glycine in *D. melanogaster* located 19 amino acids before the end of the protein. No polymorphism was found at this site in genes from a Zambian population from the Drosophila Genome Nexus data (Lack et al., 2015) nor in 13 DSPR founder strains of *D. melanogaster* (Chakraborty et al., 2019).

**Table 1.** Sites under selection in glue genes based on the Fixed Effect Likelihood Method (FEL) implemented in HyPhy. p<0.1. Sites total: Total number of codons in the alignment.

| Gene | Sites total | Sites under negative selection | Sites under positive selection |
| --- | --- | --- | --- |
| **Sgs8** | 73 | 8 | 0 |
| **Sgs7** | 74 | 11 | 0 |
| **Sgs3** | 77 | 7 | 0 |
| **Eig71Ee** | 245 | 21 | 0 |
| **Sgs5** | 144 | 29 | 0 |
| **Sgs5bis** | 142 | 16 | 0 |
| **Sgs4** | 136 | 24 | 1 |
| **Sgs1** | 250 | 24 | 0 |

### *Glue genes in D. melanogaster* show contrasting levels of genetic diversity

To gain better insight on the evolution of the eight glue genes, including the four glue genes containing difficult-to-align repeats (*Sgs1, Sgs3, Sgs4* and *Eig71Ee*), we estimated levels of polymorphism using high-quality (PacBio) genome assemblies from 15 isogenic strains of *D. melanogaster* (Chakraborty et al., 2019; Figure 4, Table 2) with the caveat that we are not sampling a natural population in equilibrium. As the evolutionary dynamics of repeat regions may be different from the rest of the sequences, population genetic statistics were estimated with and without the repeat regions for *Sgs1, Sgs3, Sgs4*, and *Eig71Ee*. The *Sgs1* gene sequence from the A7 strain was removed from the full alignment of *Sgs1* because it contained unresolved sequencing errors in the repeats. To confirm our findings, statistics were also estimated for a natural population from Zambia from the Drosophila Genome Nexus for which repeat regions of *Sgs1, Sgs3, Sgs4, Eig71Ee* contain missing data and have been removed for the analysis (Lack et al., 2015).

**Table 2.** Population genetic summary statistics of the eight glue proteins in *D. melanogaster*. Summary statistics were estimated for the fifteen *D. melanogaster* assemblies (DSPR: 13 DSPR founder lines, ORE and iso-1) as well as separately for a sampled Zambian population (ZI: from the Drosophila Genome Nexus). Chr: chromosome arm. n: number of individual genomes used for the analysis. Sites: total number of sites in the alignment (bp) and total number of sites used for the analysis in brackets. S: number of segregating sites. Singleton: number of singleton sites. Syn: number of synonymous sites. Non syn: number of nonsynonymous sites. Hd: Haplotype diversity. π: Nucleotide diversity. Θw: Watterson estimator of diversity. TajD: Tajima’s D statistic (*p<0.05, see Table S5). Values in bold indicate noticeable features discussed in the text. Genes are grouped according to their chromosome location. ^R^ Only the region containing repeats. ^1^ Sequence excluding the repeat regions.

| Gene | Pop | Chr | n | Sites | S | Singl<br>eton | Syn | Non<br>syn | Hd | $\pi$ | $\Theta_w$ | TajD |
| --- | --- | --- | --- | --- | --- | --- | --- | --- | --- | --- | --- | --- |
| **Sgs8** | DSPR | 3L | 15 | 228 [228] | 5 | 3 | 4 | 1 | 0.70 | **0.005** | 0.007 | -0.82 |
| **Sgs8** | ZI | 3L | 165 | 228 [228] | 14 | 3 | 6 | 8 | 0.92 | 0.010 | 0.012 | -0.31 |
| **Sgs7** | DSPR | 3L | 15 | 225 [225] | 1 | 0 | 0 | 1 | 0.25 | **0.001** | 0.001 | -0.40 |
| **Sgs7** | ZI | 3L | 131 | 225 [225] | 9 | 5 | 7 | 2 | 0.44 | **0.002** | 0.007 | **-1.65\*** |
| **Sgs3** | DSPR | 3L | 15 | 393 [684] | 18 | 8 | 14 | 4 | 0.72 | **0.005** | 0.008 | -1.32 |
| **Sgs3<sup>R</sup>** |  |  |  | 567 [312] | 15 | 7 | 12 | 3 | 0.48 |  |  |  |
| **Sgs3<sup>1</sup>** | DSPR | 3L | 15 | 372 [372] | 3 | 1 | 2 | 1 | 0.55 | **0.002** | 0.002 | -0.95 |
| **Sgs3<sup>1</sup>** | ZI | 3L | 81 | 823 [231] | 4 | 2 | 2 | 2 | 0.23 | **0.001** | 0.003 | -1.40 |
| **Eig71Ee** | DSPR | 3L | 15 | 1584 [1083] | 33 | 16 | 14 | 16 | 0.88 | 0.008 | 0.009 | -0.50 |
| **Eig71Ee<sup>R</sup>** |  |  |  | 699 [207] | 8 | 1 | 4 | 1 | 0.33 |  |  |  |
| **Eig71Ee<sup>1</sup>** | DSPR | 3L | 15 | 885 [876] | 25 | 15 | 10 | 15 | 0.88 | 0.007 | 0.009 | -0.86 |
| **Eig71Ee<sup>1</sup>** | ZI | 3L | 66 | 1338 [1077] | 68 | 18 | 28 | 36 | 0.95 | 0.011 | 0.013 | -0.65 |
| **Sgs5** | DSPR | 3R | 15 | 492 [494] | 7 | 0 | 4 | 3 | 0.88 | 0.007 | 0.004 | **2.10\*** |
| **Sgs5** | ZI | 3R | 197 | 492 [492] | 38 | 7 | **26** | 12 | 0.97 | 0.010 | 0.001 | **-0.86** |
| **Sgs5bis** | DSPR | 3R | 15 | 429 [429] | 7 | 2 | 5 | 2 | 0.93 | 0.007 | 0.005 | **1.15** |
| **Sgs5bis** | ZI | 3R | 196 | 429 [429] | 34 | 9 | **24** | 10 | 0.93 | 0.006 | 0.014 | **-1.58\*** |
| **Sgs4** | DSPR | X | 15 | 1074 [738] | 29 | 13 | 6 | 21 | 0.78 | 0.012 | 0.013 | 0.10 |
| **Sgs4<sup>R</sup>** |  |  |  | 633 [296] | 21 | 11 | 5 | 14 | 0.51 |  |  |  |
| **Sgs4<sup>1</sup>** | DSPR | X | 15 | 441 [441] | 7 | 2 | 2 | 5 | 0.73 | 0.006 | 0.005 | 0.86 |
| **Sgs4<sup>1</sup>** | ZI | X | 52 | 864 [552] | 45 | 25 | 15 | 27 | 1.00 | 0.011 | 0.018 | -1.40 |
| **Sgs1** | DSPR | 2L | 14 | 4437 [2943] | 243 | 150 | 68 | 179 | 0.97 | 0.019 | 0.026 | -1.42 |
| **Sgs1<sup>R</sup>** |  |  |  | 3537 [2043] | 236 | 143 | 65 | 175 | 0.97 |  |  |  |
| **Sgs1<sup>1</sup>** | DSPR | 2L | 15 | 900 [900] | 7 | 7 | 3 | 4 | 0.25 | **0.002** | 0.002 | -0.70 |
| **Sgs1<sup>1</sup>** | ZI | 2L | 87 | 3861 [996] | 74 | 29 | 28 | 44 | 0.99 | 0.009 | 0.015 | **-1.51\*** |

**Figure 4.**
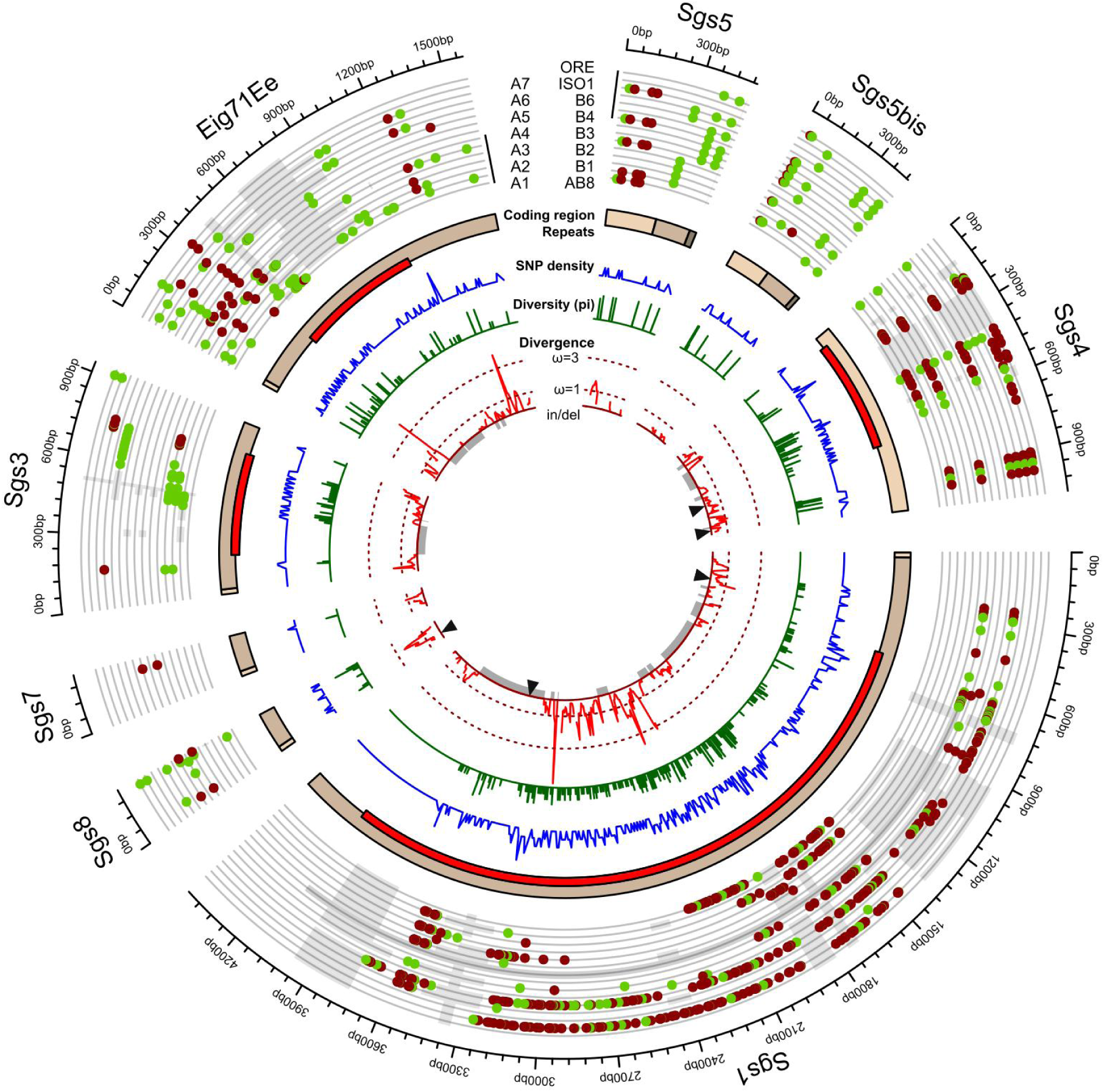
Genetic variation in glue genes across *D. melanogaster* lines and species divergence between *D. melanogaster* and *D. simulans*. We used the consensus sequence of the *D. melanogaster* multiple alignment as a reference to represent polymorphism and divergence. From outer to inner layer: sequence alignment of the 15 *D. melanogaster* lines: nonsynonymous (carmine dots), synonymous (green dots) SNPs, indels (light gray rectangles) and perfect matches (grey lines) compared to the consensus sequences, missing information for Sgs1 in A7 (grey region); coding sequence (from light to dark brown rectangles: 1st, 2nd, 3rd exons); repeat regions (red rectangles); SNP density estimated across 10 nt sliding windows (blue); π per site (green); Graph in the center: Divergence between *D. simulans* w501 and *D. melanogaster* consensus sequence: ω (red) estimated across 50 nt sliding windows (10 nt step size), deletions (grey regions) and insertions (black triangles) in *D. simulans* compared to *D. melanogaster*.

The three glue genes, *Sgs3, Sgs7*, and *Sgs8*, which are clustered on chromosome arm 3L reveal low levels of genetic diversity compared to the other glue genes (respectively, excluding repeats, π=0.002, π=0.001, and π=0.005; Table 2), *Sgs7* being the least genetically diverse with only one polymorphic site. When excluding the repeat regions, *Sgs1* also presents a low level of diversity with only seven segregating sites across over 900 base pairs (Table 2). These segregating sites precede the repeats and correspond to the doubletons found in B3 and A7 (Figure 4). Similar results were obtained with the Zambian population with *Sgs1* and *Sgs7* having a slightly significantly low Tajima’s D (respectively -1.51 and -1.65, Table 2). Altogether, these results suggest that *Sgs1, Sgs3, Sgs7*, and *Sgs8* coding sequences may have experienced recent selective sweeps.

Interestingly, *Sgs5* and *Sgs5bis* present a relatively high Tajima’s D compared to the other glue genes for the small global dataset (respectively, 2.102 and 1.146, Table 2) whereas Tajima’s D values are low when using the Zambian population, in particular for *Sgs5bi*s (TajD=-1.59, Table 2), which may suggest local adaptation or diversifying selection in certain lines from the small worldwide dataset. Interestingly, *Sgs5* shows an excess of polymorphic synonymous sites in the Zambian population (Table 2), with nonsynonymous sites being restricted to the first exon. This pattern is also observed in the 15 *D. melanogaster* lines (Figure 4). No specific polymorphism feature was detected for *Sgs4* and *Eig71Ee*.

With respect to indels, we only found two outside of the repeat regions. The first indel corresponds to a 24-bp deletion at the 3’ end of the Sgs8 sequence in the A5 strain. The second indel corresponds to a 2-bp insertion, TT, in the Sgs5 sequence of A7, located 11 bp before the end of the coding region. These two mutations cause frameshifts and change the relative position of the termination codon. These indels are not found in any other individuals (n=866) from the Drosophila Genome Nexus study, a large global survey of genetic variation in D. melanogaster (Lack et al., 2015). Future studies to test whether these two candidate mutations affect glue adhesion can involve CRISPR-directed mutagenesis.

We then examined Fst values for pairs of populations from Egypt (EG), France (FR), Raleigh (RAL) and Zambia (ZI). We found that for certain pairs of populations and for certain glue gene regions, the Fst value is among the highest 5 percentile of the distribution (Table S3, Figure S4), suggesting high differentiation of these genes in some of these populations and thus local adaptation. In particular, we found high Fst’s for Sgs4 and Sgs3 between Zambia and the three other populations.

In terms of number of repeats and repeat motifs, we found that *Sgs1* seems to vary the most between strains compared to *Sgs3, Sgs4*, and *Eig71Ee* (Figure 4, Table 2, Table S4). In particular, the *Sgs1* repeat region of ISO1 and B3 lines present only 95 and 92% sequence identity respectively with the repeat region of the consensus sequence (Figure 4). In general, higher diversity was found in the repeats than in the non-repeated sequences for all the *Sgs* genes with repeats (Figure 4), which could be explained by a lack of selection on the repeats, higher mutation rates, concerted evolution, or errors in the assemblies or alignments. We found no obvious link between adhesion force and the number of repeats with our *D. melanogaster* strains (Table S3, Spearman correlation, p>0.5, not shown).

To compare adaptive signals among glue genes in *D. melanogaster*, the ratio of nonsynonymous to synonymous substitutions (dN/dS) as well as McDonald and Kreitman (MK) tests of selection were estimated using either the 15 global *D. melanogaster* lines or the Zambian population, using *D. simulans* w^501^ as the outgroup species (Table 3). Nonsynonymous to synonymous substitutions ratio ω was also estimated using a windows approach along the nucleotides for the *D. melanogaster*/*D. simulans* pair (Figure 4). An MK test could not be performed in *Sgs7* because it contains no synonymous polymorphic sites. Repeat regions are very divergent between *D. melanogaster* and *D. simulans* and contain multiple indels (Figure 4), complicating the alignments and subsequent analyses. *Sgs8* appears to be the most diverged glue gene with a dN/dS > 1 for the species pair, *D. melanogaster*/*D. simulans*, which confirms the rapid evolution of *Sgs8* observed in Figure 3. Since dN∼dS generally denotes a neutral evolutionary process, dN/dS significantly greater than unity may indicate positive selection. The MK test was significant for *Sgs8* in the small worldwide dataset, with an excess of nonsynonymous diverged sites and DoS=0.55 (Table 3), but not in the Zambian population (Table 2). The MK test was also slightly significant for *Sgs5* and *Sgs5bis* and a DoS>0 in the Zambian population (Table 3).

**Table 3.** Glue gene divergence between *D. melanogaster* and *D. simulans*. Summary statistics were estimated for the fifteen *D. melanogaster* assemblies (DSPR: 13 DSPR founder lines, ORE and iso-1) in addition to a Zambian population from the Drosophila Genome Nexus (ZI). Chr: c=Chromosome arm. n: Number of *D. melanogaster* individual genomes used for the analysis. Codons: Total number of codons in the alignments and total number of codons used for the analysis in brackets. Dn: Number of nonsynonymous divergent sites. Ds: Number of synonymous divergent sites. Pn: Number of nonsynonymous polymorphic sites. Ps: Number of synonymous polymorphic sites. MK G: McDonald and Kreitman test’s G value. MK p: p-value of the G-test without multiple corrections. DoS: Direction of selection. Values in bold indicate particular features discussed in the text. For *Sgs1, Sgs3, Sgs4*, and *Eig71Ee*, statistics were calculated on sequences excluding the repeat regions.

| Gene | Pop | Chr | n | Codons | Dn | Ds | Pn | Ps | MK G | MK p | DoS | dN/dS |
| --- | --- | --- | --- | --- | --- | --- | --- | --- | --- | --- | --- | --- |
| ***Sgs8*** | DSPR | 3L | 15 | 75 [73] | **24** | 8 | 1 | 4 | **5.63** | **0.018** | **0.55** | **1.06** |
| ***Sgs8*** | ZI | 3L | 165 | 75 [75] | 22 | 8 | 8 | 6 | 1.13 | 0.29 | 0.16 |  |
| ***Sgs7*** | DSPR | 3L | 15 | 74 [74] | 7 | 8 | 1 | 0 | NA | NA | -0.53 | 0.29 |
| ***Sgs7*** | ZI | 3L | 131 | 74 [74] | 7 | 8 | 2 | 7 | 1.49 | 0.22 | 0.24 |  |
| ***Sgs3*** | DSPR | 3L | 15 | 123 [119] | 39 | 33 | 1 | 2 | 0.51 | 0.48 | 0.21 | 0.39 |
| ***Sgs3*** | ZI | 3L | 81 | 76 [76] | 11 | 11 | 2 | 2 | 0.00 | 1.00 | 0.00 |  |
| ***Eig71Ee*** | DSPR | 3L | 15 | 294 [250] | 41 | 17 | 12 | 9 | 1.25 | 0.26 | 0.14 | 0.73 |
| ***Eig71Ee*** | ZI | 3L | 66 | 358 [142] | 40 | 21 | 24 | 25 | 3.08 | 0.08 | 0.17 |  |
| ***Sgs5*** | ZI | 3R | 197 | 163 [163] | 11 | 6 | 13 | 27 | **5.07** | **0.02** | **0.32** |  |
| ***Sgs5bis*** | DSPR | 3R | 15 | 142 [142] | 8 | 4 | 2 | 5 | 2.64 | 0.11 | 0.38 | 0.41 |
| ***Sgs5bis*** | ZI | 3R | 196 | 142 [142] | 8 | 3 | 10 | 24 | **6.49** | **0.01** | **0.43** |  |
| ***Sgs4*** | DSPR | X | 15 | 146 [139] | 24 | 23 | 5 | 2 | 1.05 | 0.31 | -0.20 | 0.38 |
| ***Sgs4*** | ZI | X | 52 | 183 [162] | 35 | 25 | 24 | 15 | 0.10 | 0.75 | -0.03 |  |
| ***Sgs1*** | DSPR | 2L | 15 | 299 [281] | 41 | 40 | 3 | 2 | 0.17 | 0.68 | -0.094 | 0.32 |
| ***Sgs1*** | ZI | 2L | 87 | 331 [294] | 38 | 30 | 42 | 25 | 0.65 | 0.85 | -0.07 |  |

## Discussion

We provide, for the first time, global estimates of phenotypic variation of pupal adhesion in an insect species. Our adhesion assay is based on measuring adhesion of individual pupa from twelve isogenized *D. melanogaster* lines originating from different global locations. Variation between individuals is high and cannot be explained solely by measurement errors since we used a 5N force sensor with a precision of ±0.5% on the read value. The high variance can be explained by different environmental, developmental, physiological, behavioral, and morphological parameters: for example due tobut individual variation in the position or the shape of the pupa, the way glue spreads over the substrate, or to stochastic variation in glue production, in glue composition, or in very local conditions at the moment when the glue was excreted. It could also be due to variation in the position of the initial crack in the glue, which leads to detachment (Borne et al., 2020; Borne et al., 2021). Nonetheless, we found that 26% of the variation is explained by strain and could thus reflect adaptation of the glue to the natural climatic conditions experienced by flies. In particular, we identified two lowly adhesive lines, A5 (Athens, Greece) and B4 (Riverside, CA, USA), and two strongly adhesive lines, A7 (Ken-ting, Taiwan) and B6 (Ica, Peru). While A5 and B4 lines are from locations with similar climates (hot summer, little rain in winter), A7 (hot summer, heavy rain in winter) and B6 (hot summer, warm and dry winter) lines are not. To our knowledge, the impact of climate on glue strength has never been studied so far. As glue protects from predation (Borne et al., 2021), glue could also be adapted to the local predation pressure. More lines should be tested to examine possible correlations between adhesion force and specific environmental factors. Yet despite the observed large variance among *D. melanogaster* lines, we did not find a significant difference in adhesion between *D. melanogaster* and other *D. melanogaster* subgroup species tested except for *D. mauritiana*. Only one or two strains have been tested per species which is not enough to appreciate the intraspecific variation for those species. Furthermore, species and lines may vary in other phenotypes related to glue adhesion force (substrate specificity, plasticity to temperature or humidity conditions, etc). The glue proteins have mucin-like regions and some of them may thus have antimicrobial properties like some mucins, i.e., it is possible that the glue of various populations and species is adapted to resist different pathogens.

The glue is produced by the salivary glands and glue production is the only known function of these glands in *D. melanogaster* during the third larval wandering stage (Beňová-Liszeková et al., 2021). We found that the glue genes are among the most highly expressed genes in the salivary glands at the wandering larval stage. Other small uncharacterised genes also display the same tissue-specific, stage-specific, and high expression levels: they represent interesting candidate proteins that may also be involved in *D. melanogaster* glue composition. Glue genes evolve rapidly across Drosophila species (Da Lage et al., 2019). Our work suggests that transcriptomics might be an effective method to identify candidate glue genes in other Drosophila species. To our knowledge, no transcriptome has been done for larval salivary glands in Drosophila species outside of *D. melanogaster*.

The degree of codon bias is usually positively correlated with the level of expression (Sharp and Li, 1987). Surprisingly, in the case of the salivary glands of third instar wandering larvae, we find two groups of highly expressed genes: a first one with high CAI (encoding for ribosomal proteins, cytoskeleton components, etc.) and another one with low to medium CAI and thus little codon bias (glue genes and genes of unknown function) (Table S1). Besides protein function, the DNA sequence of the glue genes may be constrained not only by translation processes and tRNA genes, but also by factors specific to the salivary gland or the glue genes, such as DNA stability due to the presence of numerous DNA copies within polytene chromosomes and puffs where mRNA synthesis occurs (Zhimulev et al., 2004). Another explanation is that the coding sequence of the glue genes has evolved more rapidly than other genes (Da Lage et al., 2019) and that there has not been enough time for codon bias to be optimized. Interestingly, recent studies suggest that codon usage could also impact protein structure (Orešič and Shalloway, 1998; Pechmann and Frydman, 2013; Zhou et al., 2015, 2009 reviewed in Liu, 2020). As protein folding occurs simultaneously during translation, it can be affected by translation speed. In particular, several studies suggest that unstructured domains tend to be less codon-biased than structured ones (Zhou et al., 2015), which could explain an enrichment in rare codons in the disordered glue genes Sgs1, Sgs3, Sgs4 and Eig71Ee. Additionally, rare codons could enhance membrane targeting and secretion efficacy of secreted proteins (Pechmann et al., 2014).

Using high-quality PacBio genome assemblies, we annotated the glue genes of the sister species *D. simulans, D. mauritiana, D. sechellia* and the more distant species *D. yakuba* and *D. teissieri*, based on BLAST and synteny. We identified duplicates of *Sgs3* in *D. yakuba* and *D. teissieri* which were not found using short-read based assemblies (Da Lage et al., 2019). The discovery of these new duplicates indicates that high quality genomes are necessary to precisely survey the evolution of such rapidly evolving genes. However, alignment in the repeat regions remains challenging as the repeats are very divergent. In accordance with Da Lage et al., 2019, we found that *Sgs8*, similar in sequence identity to *Sgs7*, was the most rapidly evolving gene among the eight glue genes in *D. melanogaster*, potentially due to post-duplication evolutionary dynamics (e.g., Lynch and Conery 2000). However, the *Sgs8* sequence is relatively short and divergence observed in this gene could simply be due to sampling bias and thus be the result of non-adaptive evolution. Contrary to what was observed in Da Lage et al. (2019), we did not find high levels of divergence for *Sgs1* between *D. melanogaster* and *D. simulans*. This previous study used the full *Sgs1* coding region including repeats and it is likely that the estimation was biased by the repeat regions (dN/dS ratio value obtained with the Yang and Nielsen, 2000 method with repeats: 1.4, without repeats: 0.5).

Furthermore, we observed rapid evolution of *Sgs3* coding sequence in the *D. yakuba* branch after the duplication event, which occurred before the *D. yakuba* - *D. teissieri* divergence. This rapid evolution could reflect specific and recent adaptation of the glue in *D. yakuba*. In agreement with our findings, Da Lage et al. (2021) previously reported accelerated evolution of *Sgs3* in the *D. yakuba*/*D*.*santomea* branch. Using Fixed Effect Likelihood methods, we identified multiple sites under purifying selection in the eight glue genes and only one site under positive selection in *Sgs4*. The presence of sites under purifying selection highlights the possible important functions of the glue genes. So far, functions of the glue genes have not been tested and glue proteins do not contain known functional domains (Farkas, 2016; Da Lage et al., 2019). Identified sites should be further investigated and could lead to a better understanding of the mechanism of action of the glue genes.

We examined genetic variation in the glue genes within the 12 *D. melanogaster* phenotyped lines and 3 others and compared our results with a true population from Zambia. We found that *Sgs1, Sgs3, Sgs7*, and *Sgs8* contain a low level of diversity in their non-repeat regions, suggesting that they may have experienced recent selective sweeps. In contrast, we found high diversity in the repeats, in agreement with Da Lage et al., 2019, suggesting that the repeats have the capacity to evolve rapidly due to lower constraints in the sequence. With respect to *Sgs5* and *Sgs5bis*, the analyses based on the fifteen *D. melanogaster* lines and the Zambian population present contrasting Tajima’s D estimates, which may be explained by recent local adaptations of the smaller worldwide dataset. The MK test was slightly significant in *Sgs5* and *Sgs5bis* when using the Zambian population, suggesting potential adaptive evolution of these genes.

Mapping phenotype to genotype still remains an elusive challenge. In this study, we surveyed the genomic landscape to identify the genetic variants and evolutionary processes that may have played a role in the widespread species-level variation of pupal adhesion. Understanding the genetic basis of pupal adhesion in Drosophila provides an excellent model as it combines simplicity at the phenotypic level--a well-defined phenotypic trait expressed at a precise developmental stage in a single tissue--with complexity at the genomic level, i.e., multiple genes with either long coding regions and large repeats or short coding regions. This study provides the foundation for future work that can directly connect these variants with fitness during this particularly precarious stage of development.

## Methods

### Fly samples

Flies were cultured at 25 °C in plastic vials on standard medium [4 liters: 83.5 g yeast, 335.0 g cornmeal, 40.0 g agar, 233.5 g saccharose, 67.0 ml Moldex, 6.0 ml propionic acid]. We used all 13 DSPR founder lines for which their genome has been PacBio-assembled (Chakraborty et al., 2019, gift from Antony Long) except AB8, B1, and A3 lines because these strains are no longer available. We also used ORE (Oregon R, Bloomington Stock Center #25211) and iso-1 (provided by Jean-Luc Da Lage) *D. melanogaster* strains, *D. simulans* w[501] and md221 strains, *D. mauritania* mau12, *D. yakuba* NY73PB, and Tai18E2 (all 5 strains provided by Peter Andolfatto).

### Estimating adhesion strength

Pupae attached to glass slides (Menzel Superfrost microscope glass slide from ThermoScientific™ #AGAB000080) were prepared as previously described (Borne et al., 2021, 2020). Adhesion force of an individual pupa was measured by detaching individual pupa from the glass slides using a universal test machine (LS1S/H/230V Lloyd Instruments, Ametek®) with a 5N force sensor (YLC-0005-A1 Lloyd Instruments™, Ametek®) as previously described (Borne et al., 2021). Adhesion assays were performed at ambient room temperature and humidity. Force-time curves were recorded using NEXYGENPlus software (Lloyd Instruments™). The adhesion force of each individual corresponds to the maximal force reached during the experiment. Pupae which did not detach from the glass slide (representing less than 10% of the tested pupae) and pupae whose pupal case broke during the pulling phase (representing less than 11% of the tested pupae) were not taken into account for further analysis (see Dataset1.ods in Dryad). Images of glue prints remaining on the glass slides after detachment were taken with a Keyence VHX2000 microscope with a VH-Z20R/W objective x100. Images were anonymized and print areas were measured manually using imageJ (1.50d, java 1.8.0_212, 64-bit). We measured the contact area between the glue and the substrate as described previously, corresponding to the area on which the pupa was in contact with the substrate (Borne et al., 2020; Borne et al., 2021). Prints for which there was no glue left on the slides (n=88) or which were damaged after the tests (lost (n=5), wet (n=6) or dirty slides (n=2)) were not included in the analysis.

Differences in pull-off force between *D. melanogaster* strains and between species were tested by one-way ANOVA using the aov() R function followed by multiple pairwise comparison tests using the Tukey test with the TukeyHSD() R function. To compare the different species, we conducted the ANOVA using species as groups. The 12 lines of D. melanogaster were placed in the same group. Functions belong to the stats core package of R (v 3.6.3) (https://www.r-project.org/). The proportion of the phenotypic variance explained by strain was measured by dividing the sum square of deviation of the mean of treatments (i.e. strains) by the total sum square obtained by ANOVA. To test whether adhesion forces were correlated with glue-substrate contact areas in *D. melanogaster*, we performed a standardized major axis (sma) regression using the sma() function from the rsmartr R package (Warton et al., 2012).

### Analysis of transcriptome data and codon bias

We uploaded the RPKM values of the genes with expression levels “moderately high” and above in salivary gland wandering larvae based on modENCODE high-throughput RNA-seq (Graveley et al., 2011) using the Flybase interface (https://flybase.org/rnaseq/profile_search). To calculate the fold expression difference between glue genes and median genes, we considered as median the genes with “moderate” expression levels which represent percentiles 51-75 of RPKM values (Flybase reports: https://flybase.org/reports/FBrf0221009.html). To collect the CDS sequences corresponding to these genes, we downloaded the genome of *D. melanogaster* iso1 from Flybase (release r6.39) with the corresponding gtf file. We extracted all the CDS with gffread (v0.9.12; Pertea and Pertea, 2020), filtering for each gene’s longest CDS using a Python script provided by Mathilde Paris. The codon adaptation index (CAI) (Sharp and Li, 1987) were estimated on CDS using CAIcal (v1.4; (Puigbò et al., 2008) and a codon usage reference table of *D. melanogaster* (Nakamura et al., 2000). Codon usage of the glue genes for threonine and proline was performed on the A1 strain coding sequences using codonw (Peden, 1999), http://codonw.sourceforge.net/).

### Assemblies used for genomic analyses

*Due to the relatively high repetitive nature of the glue gene sequences, we used fifteen high-quality D. melanogaster* genomic assemblies including the reference genome iso-1 (Dmel R6), recently published PacBio assemblies from 13 founder lines of the Drosophila Synthetic Population Resource (DSPR) and the Oregon R strain (Chakraborty et al., 2019, see Data availability). Genome assemblies of *D. mauritiana* (mau12) and *D. sechellia* (sech25) were downloaded from NCBI (see Data availability). PacBio genome assemblies of *D. yakuba* (NY73PB), *D. teissieri* (GT53w) and *D. simulans* (w^501^) were kindly provided by Peter Andolfatto (see Data availability).

### Manual curation of gene sequences from PacBio genomic alignments

Glue gene coding sequences were identified in each genome assembly using BLASTN implemented in Geneious prime (v.2019.1.3) and using previous glue gene models as queries (Da Lage et al., 2019). Coding regions were annotated manually using Geneious prime (v. 2019.1.3). A previously unidentified duplication of the *Sgs3* gene was found in *D. yakuba* and *D. teissieri* assemblies. Please note that the *Sgs1* locus is inverted in the *D. yakuba* strain NY73PB and that the *Sgs5*-*Sgs5bis* locus is inverted in *D. melanogaster* A2 strain as already mentioned in King et al., 2012. For Figure S3, the nucleotide alignment was performed in Geneious prime (v. 2019.1.3) using Muscle (v3.8.425) and the dot plot was done in Geneious prime (v. 2019.1.3) with a word size of 15 nucleotides.

When a frameshift or premature stop codon was detected in an individual *D. melanogaster* line, we blasted Illumina raw read data from the same strain (King et al., 2012, see Data availability) in NCBI’s SRA using no filter for low complexity regions. If the reads did not reveal a frameshift or premature stop codon, we treated the mutation as an error and corrected it. Individual assemblies were thus corrected for the B1 strain (coding sequence of *Sgs4*, chr X:3067004, replaced T by A), AB8 strain (coding sequence of *Sgs3*, chr 3L:11420384-11420385, T was added), and A1 strain (coding sequence of *Sgs3*: 3L:11445654-11445655, TAAGCCCA was added; 3L:11445657 C replaced by A; 3L:11445715-11445716, G was added; 3L:11445711-11445712 CCA was added). The coding sequence of *Sgs1* from the A7 strain was not used in the full alignment of *Sgs1* sequences because it probably contains multiple errors in the repeat regions that we could not correct with confidence. For the *D. teissieri* assembly, one deleted nucleotide in *Sgs1* created a frameshift at amino acid position 244. Since we did not have access to its raw Illumina data, we manually annotated this site with a ‘?’ (2L:5055203-5055204).

### Interspecific divergence of glue genes

To estimate the evolutionary rate of each glue gene across species, we aligned the coding sequences of *D. melanogaster* (A1), *D. simulans* (w501), *D. mauritiana* (mau12), *D. sechellia* (sec25), *D. yakuba* (NY73PB) and *D. teissieri* (GT53w) using MUSCLE implemented in MEGA-X v.10.1.7 (Kumar et al., 2018). Codon-guided alignments were corrected manually if needed. Repeated parts of *Sgs1, Sgs3, Sgs4* and *Eig71Ee* genes and indels were removed manually. The beginning and the end of the repeat regions were determined visually according to the protein sequences. For *D. teissieri* and *D. yakuba*, which contain multiple copies of *Sgs3* and *Sgs7*, we only use one copy for the *Sgs7* alignment as the two copies were almost identical (see Results) and we used all copies for *Sgs3* alignments. Nonsynonymous (dN) substitution rates were determined for each lineage using a one ratio model (M0 model) and a free ratio model codon with the frequency model F3×4 in CODEML from the PAML package (v4.7, Yang, 1997). The one ratio model (NSsites=0, model=0) allows one average ω for all branches whereas the free ratio model (NSsites=0, model=1) allows different ω for each branch of the tree. For all genes except *Sgs3*, we used the unrooted species tree (((simulans, mauritiana, sechellia), melanogaster), yakuba, teissieri). For *Sgs3*, we used the gene tree obtained with PhyML plugin in Geneious prime (v. 2019.1.3) with the K80 substitution model and bootstrap 100 (v3.3.20180621, Guindon et al., 2010): ((sim_Sgs3, mau_Sgs3, sec_Sgs3),mel_Sgs3), (yak_Sgs3, tei_Sgs3), (yak_Sgs3bis, (tei_Sgs3bis, tei_Sgs3ter))); (Figure S2). Statistical significance between the two PAML models was determined by likelihood ratio tests assuming a chi-square distribution. Since the free ratio model was not significantly better than the one ratio model except for *Sgs3* (Table S2), dN values from the free ratio model were used for *Sgs3* and values from the one ratio model were used for the other glue genes. dN trees were drawn in ngphylogeny.fr using the Newick display tool (v.1.6; (Junier and Zdobnov, 2010) and modified on iTOL (Letunic and Bork, 2021).

### Identifying sites under selection

To identify sites under selection, we applied the Fixed Effects Likelihood (FEL) methods (Kosakovsky Pond and Frost, 2005) implemented in HyPhy (Kosakovsky Pond et al., 2020) on the webportal http://datamonkey.org (Weaver et al., 2018) to the multiple sequence alignment used for the analysis of evolutionary rate, without the *Sgs3* and *Sgs7* duplicates. Since *Sgs3, Sgs7*, and *Sgs8* are close to each other (<5 kbp) on chromosome 3L, we concatenated the coding sequences of these three genes to increase the power of the analysis. Similarly, we concatenated the coding sequences of the tandem duplicates *Sgs5* and *Sgs5bis* (distance<1.7 kbp).

### Analyses of natural populations of *D. melanogaster*

To check that the results obtained from the fifteen geographically diverse *D. melanogaster* strains were similar to those from a natural population, we estimated the same population genetic parameters using the aligned sequences of the Zambia (ZI) population from the Drosophila Genome Nexus (Lack et al., 2015), a collection of over 1,000 Drosophila genomes spanning dozens of global populations. Zambia is proximal to the ancestral origins of this now cosmopolitan species. We downloaded the sequences from Popfly (Hervas et al., 2017; https://popfly.uab.cat/), annotated and extracted the coding sequences of *Sgs1, Sgs3, Sgs4, Sgs5, Sgs5bis, Sgs7, Sgs8* and *Eig71Ee* using Geneious prime (v. 2019.1.3). Individuals with missing data or with a premature stop codon in the sequences of interest were not used. For *Sgs1, Sgs3, Sgs4* and *Eig71Ee*, we removed the part of the sequences containing repeats since they contained missing data in all of the individuals, corresponding, respectively, to positions 573-3437, 175-867, 150-461 and 679-393 of the coding regions. To estimate divergence, we used the aligned *D. simulans* sequence provided in the Drosophila Genome Nexus (Stanley Jr and Kulathinal, 2016). We retrieved the 1kb-non-overlapping-window Fst values available on the Popfly website (https://popfly.uab.cat/files/?dir=fst) for all the possible pairs of the following Drosophila Genome Nexus populations: Egypt (EG), France (FR), Raleigh (RAL) and Zambia (ZI). Within the distribution of all the Fst values estimated across the whole genome for a given pair of populations, the position of the Fst value obtained for genome windows containing the glue gene coding sequences was calculated with a custom R script.

### Analysis of polymorphism and McDonald-Kreitman test

Coding sequences for each glue gene were aligned across the 15 *D. melanogaster* lines using MUSCLE and corrected manually as above. Population genetics parameters were estimated with and without the repeats for *Sgs1, Sgs3, Sgs4*, and *Eig71Ee*.

To quantify genetic variation in *D. melanogaster*, we estimated the number of segregating sites (S), the number of synonymous and nonsynonymous sites, nucleotide diversity (π) (Nei and Li, 1979), Watterson’s θ (θw) (Watterson, 1975), haplotype diversity (Hd) and Tajima’s D (TajD) (Tajima, 1989) in DNAsp v.6.12.03 x64 (Rozas et al., 2017). Significance of Tajima’s D test was obtained by comparing the observed values with values from 10000 neutral coalescent simulations generated by DNAsp with no recombination and by fixing the number of segregating sites (Table S5). vcftools (v 0.1.16) (Danecek et al., 2011) was applied to calculate sliding windows of π and DNAsp (v.6.12.03 x64) was used to estimate SNP density and ω sliding windows. ω sliding windows were calculated using the consensus sequence of *D. melanogaster* obtained from the fifteen genomes alignment and the sequence of *D. simulans* (w^501^), *D. mauritiana* (mau12) or *D. sechellia* (sech25) (Figure S5).

Using *D. simulans* w^501^ as an outgroup, we performed a McDonald-Kreitman test (McDonald and Kreitman, 1991) and estimated the direction of selection (Stoletzki and Eyre-Walker, 2011). We also estimated the ratio of nonsynonymous to synonymous divergence dN/dS across species (Nei and Gojobori, 1986).

### Analysis of repeats

To count the number of repeated motifs in Sgs1, Sgs3, Sgs4, and Eig71Ee proteins, we first translated the coding sequences of the glue genes of the fifteen *D. melanogaster* assemblies in Geneious prime (v. 2019.1.3), determined the motif sequence based on (Da Lage et al., 2019)) and visually annotated the beginning and the end of the repeat regions. We then divided the length of the regions by the length of the motif and rounded the number of repeats obtained to the closest integer.

## DATA availability

GenBank assembly accession for iso-1, mau12, sech25, NY73PB, GT53w and W^501^: GCA_000001215.4, GCA_004382145.1, GCA_004382195.1, GCA_016746335.2, GCA_016746235.1, GCA_016746395.1

Assemblies of DSPR founder lines and Oregon R: PRJNA418342

Illumina data of the DSPR founder lines and Oregon R: SRA051316

Raw data, alignments and scripts have been deposited in the Dryad Digital Repository, doi:10.5061/dryad.v6wwpzgwk, temporary link for the review process: https://datadryad.org/stash/share/KN532B9hnGG_v17jpC2ZqyM4UQSnxK8GS1BkaZgp3oc).

## Acknowledgements

We thank Peter Andolfatto for sharing fly stocks and early access to genome assemblies, Tony Long and Jean-Luc Da Lage for sharing fly stocks, Michael Lang for his help with phylogenetic trees, and Aurelie Surtel for drawing the outlines of all the glue prints. We also thank the Bloomington Drosophila Stock Center (NIH P40OD018537) for fly stocks and Mathilde Paris for sharing her script. The research leading to this paper has received fundings from CNRS as part of the MITI interdisciplinary action “Défi Adaptation du vivant à son environnement” 2020 and from the European Research Council under the European Community’s Seventh Framework Program (FP7/2007-2013 Grant Agreement no. 337579) to V.C.-O. FB was supported by a PhD fellowship from Ecole normale superieure Paris-Saclay. RJK’s sabbatical visit to V.C.-O’s laboratory was partly supported by the LabEx “Who Am I?” #ANR-11-LABX-0071 and the Université de Paris IdEx (ANR-18-IDEX-0001) funded by the French Government through its “Investments for the Future” program.

