## Supplemental Figures and Tables for "Glue genes are subjected to diverse selective forces during Drosophila development"

### Supplementary Tables and Figures

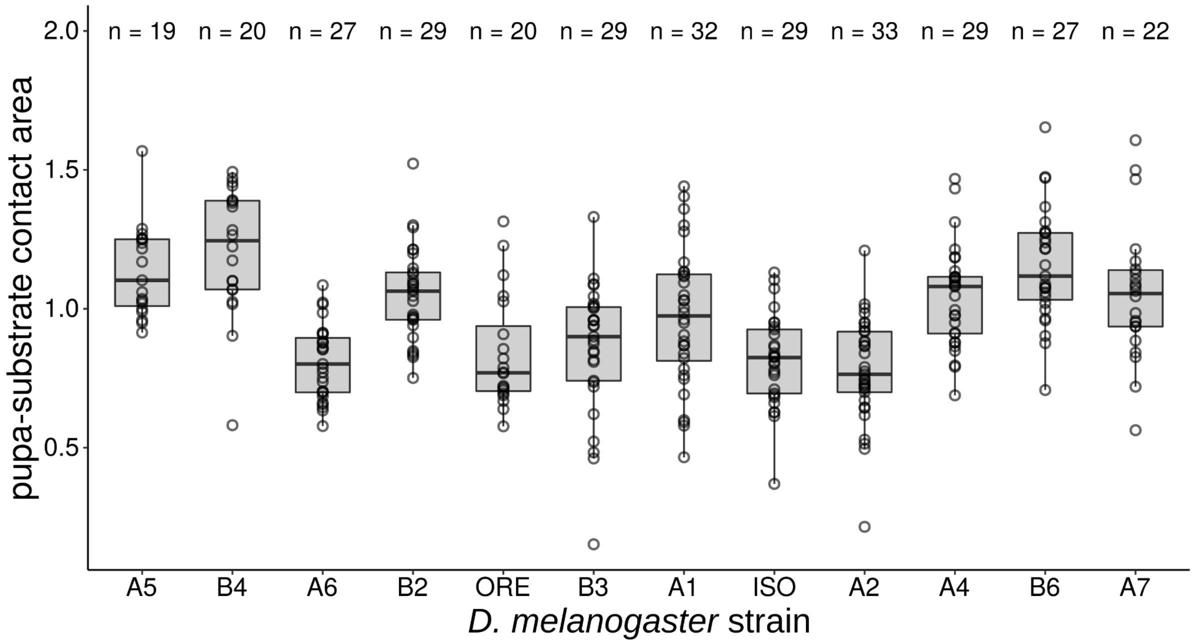


**Figure S1. Pupa - substrate contact area of the 15 lines of *D. melanogaster*.** Pupa - substrate contact area indicates the area of contact in $mm^2$ between the pupa and the substrate measured on the prints left by the glue after pupa detachment. Each dot corresponds to a single pupa and n indicates the total number of measured pupae for each line. Boxplots are defined as previously (Fig 4). Two lines are significantly different from each other when they do not share a letter (ANOVA followed by all pairwise comparisons after Tukey correction).

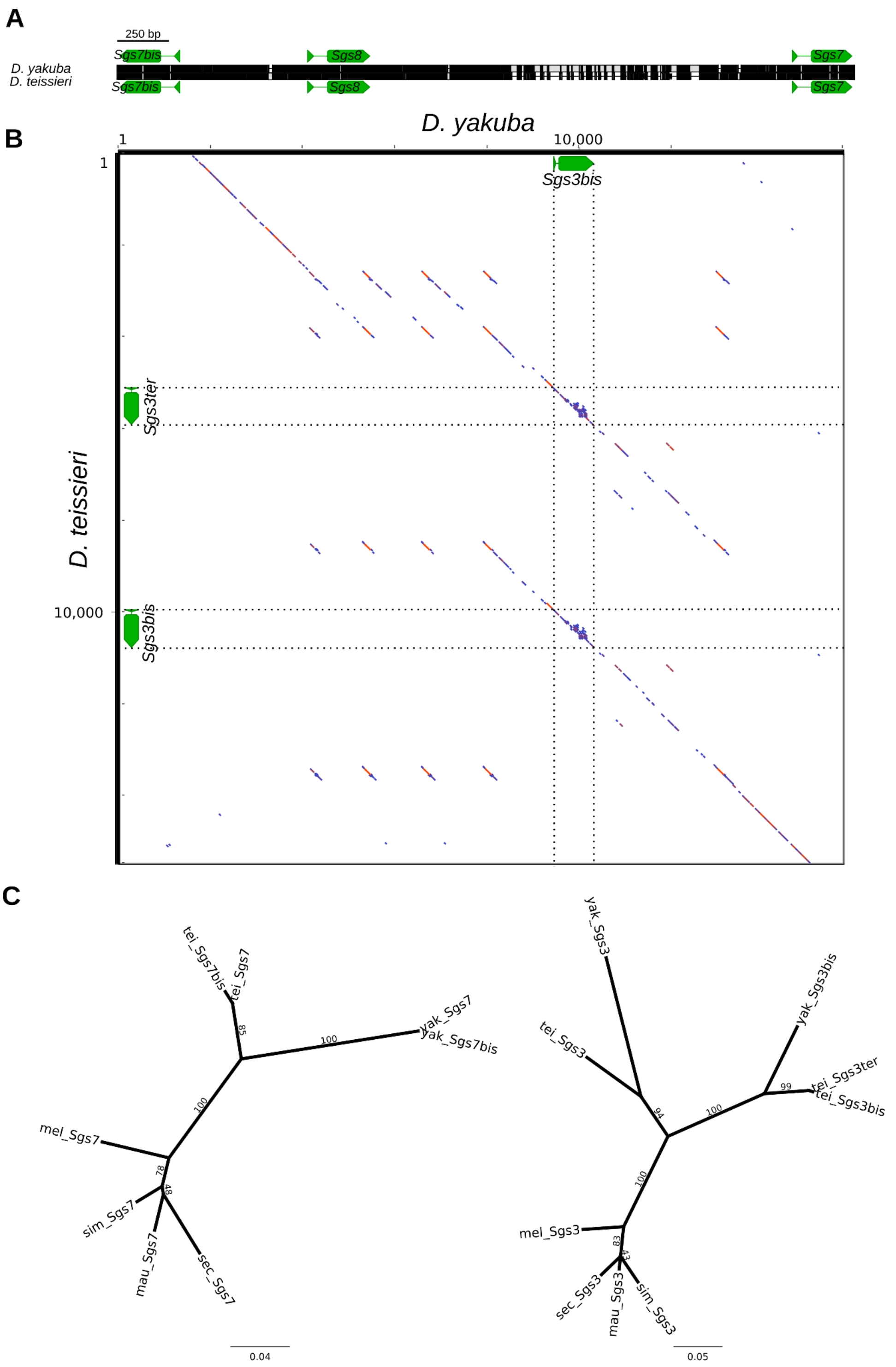


**Figure S2. *Sgs3* and *Sgs7* duplicates in *D. yakuba* and *D. teissieri*.** A) Alignments of *Sgs3bis* and *Sgs7* genomic regions between *D. yakuba* and *D. teissieri*. The nucleotide alignment is coloured in black when the sequences are identical or in white when not. B) Dotplot between *D. yakuba* and *D. teissieri Sgs3bis* genomic regions. C) Gene tree of *Sgs7* and *Sgs3* (PhyML plugin in Geneious, K80 model, bootstrap 100). Numbers on the tree branches indicate the number of bootstrap replicates which support each branch. Scale represents substitution rate.

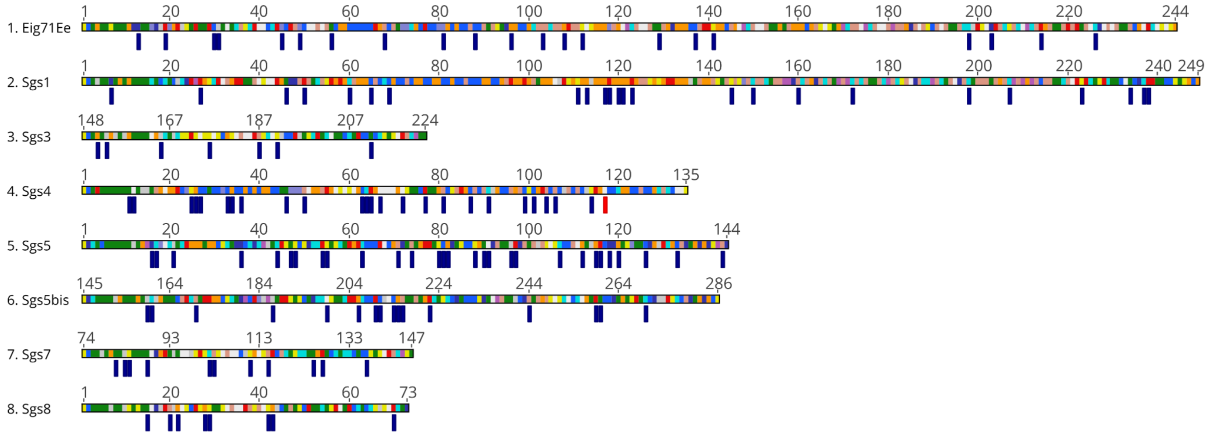


**Figure S3. Sites under selection.** Sites identified as under selection according to our PAML analysis are presented on the *D. melanogaster* glue protein sequences. Numbers indicate amino acid positions. Blue ticks indicate sites under negative selection and the red tick points to the unique sites under positive selection (in *Sgs4*).

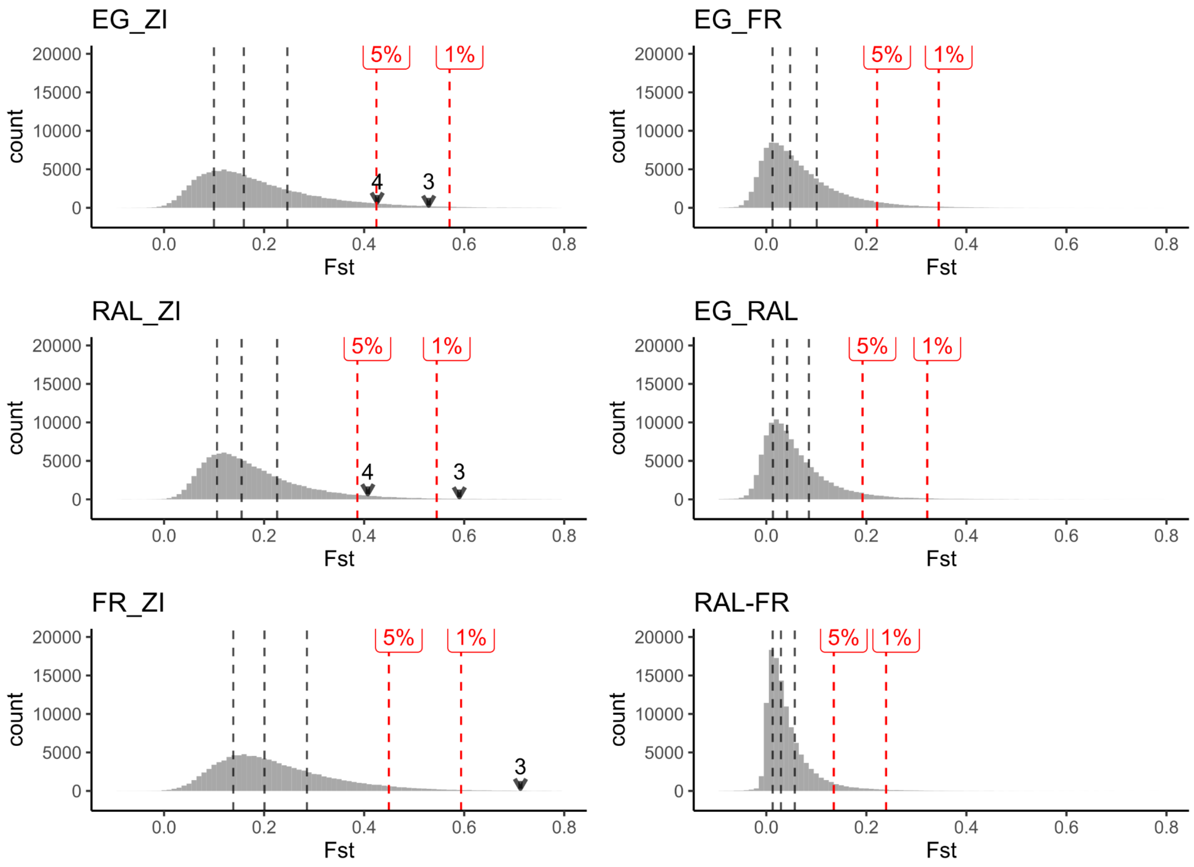


**Figure S4. Distribution of Fst estimates (1-kbp windows) across the entire genome for different pairs of populations. EG: Egypt, ZI: Zambia, FR: France, RAL: Raleigh.** The three black dashed lines correspond to the first, second (median) and third quartile of the distribution. The red dashed lines correspond to the limit of the 5% and 1% top Fst estimates. Arrows correspond to the glue genes (3: Sgs3, 4: Sgs4) which are within the top 5% Fst.

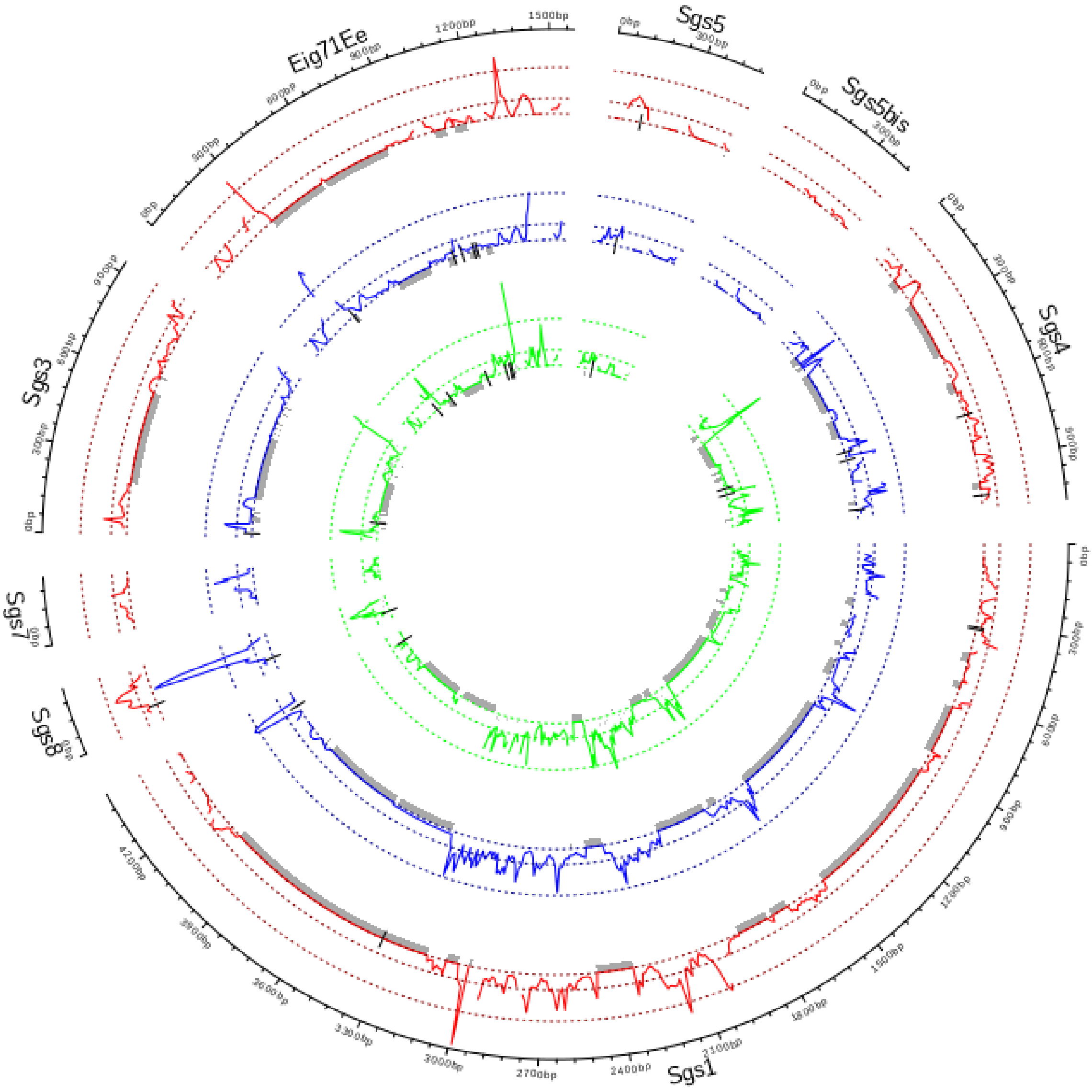


**Figure S5. Circos plot showing divergence between *D. melanogaster* and its sister species.** ω 50 nt sliding windows (10 nt step size) is shown for the following pairs: *D. simulans* / *D. melanogaster* (red), *D. sechellia* / *D. melanogaster* (blue), *D. mauritiana* / *D. melanogaster* (green). Alignments were done using *D. melanogaster* consensus sequence of the 15 lines alignment. Deletions compared to *D. melanogaster* are indicated by grey rectangles and insertions by black lines.

| FBgn | Gene Symbol | RPKM Value | CAI | Gene Information |
|---|---|---|---|---|
| FBgn0003374 | *Sgs4* | 340470 | 0.695 | glue gene |
| FBgn0003377 | *Sgs7* | 155160 | 0.689 | glue gene |
| FBgn0003373 | *Sgs3* | 90995 | 0.748 | glue gene |
| FBgn0004592 | *Eig71Ee* | 70922 | 0.63 | glue gene |
| FBgn0003375 | *Sgs5* | 38596 | 0.701 | glue gene |
| FBgn0003378 | *Sgs8* | 22325 | 0.691 | glue gene |
| FBgn0032400 | *CG6770* | 12717 | 0.934 | predicted to be involved in negative regulation of cell cycle; negative regulation of cell population proliferation; and regulation of transcription by RNA polymerase II |
| FBgn0038523 | *Sgs5bis* | 11931 | 0.759 | glue gene |
| FBgn0003372 | *Sgs1* | 11727 | 0.609 | glue gene |
| FBgn0034901 | *CG11300* | 7449 | 0.639 | unknown function |
| FBgn0051698 | *CG31698* | 6489 | 0.67 | unknown function |
| FBgn0052198 | *CG32198* | 5208 | 0.639 | unknown function |
| FBgn0066084 | *RpL41* | 4976 | 0.758 | ribosomal protein L41 |
| FBgn0040565 | *CG7606* | 4695 | 0.689 | unknown function, clustered with *Sgs5-Sgs5bis* |
| FBgn0031512 | *CG15404* | 3549 | 0.743 | unknown function |
| FBgn0034899 | *CG13560* | 3323 | 0.745 | unknown function |
| FBgn0259950 | *CG42460* | 3204 | 0.599 | predicted to enable serine-type endopeptidase inhibitor activity. |
| FBgn0036467 | *CG12310* | 2910 | 0.719 | unknown function |
| FBgn0085195 | *CG34166* | 2885 | 0.712 | unknown function |
| FBgn0263762 | *CG43679* | 2237 | 0.623 | unknown function |
| FBgn0026372 | *RpL23A* | 2023 | 0.831 | ribosomal protein L23A |
| FBgn0000042 | *Act5C* | 1885 | 0.847 | actin 5C |
| FBgn0284245 | *eEF1alpha1* | 1863 | 0.841 | eukaryotic translation elongation factor 1 alpha 1 |

| | | | | |
|---|---|---|---|---|
| FBgn0034138 | *RpS15* | 1823 | 0.851 | ribosomal protein S15 |
| FBgn0026084 | *cib* | 1759 | 0.878 | actin binding protein involved in brain development and the remodeling of the larval central nervous system |
| FBgn0003514 | *sqh* | 1546 | 0.845 | regulatory light chain of the nonmuscle type 2 myosin |
| FBgn0003941 | *RpL40* | 1514 | 0.877 | ribosomal protein L40 |
| FBgn0040393 | *CG14265* | 1438 | 0.787 | unknown function |
| FBgn0051496 | *CG31496* | 1408 | 0.604 | involved in multicellular organism reproduction |
| FBgn0285952 | *eEF5* | 1403 | 0.711 | eukaryotic translation elongation factor 5 |

**Table S1. List of the 30 most highly expressed genes in the salivary glands at the L3 wandering stage in *D. melanogaster* (adapted from Dataset1.ods in Dryad).** Gene names and coding sequences used to calculate CAI are based on genome assembly Dmel_R6.39. RPKM values are from Graveley et al., 2011. Gene information was manually retrieved from FlyBase (http://flybase.org/) for each gene.

| gene | One ratio model | Free ratio model | df | P-value |
|---|---|---|---|---|
| ***Sgs8*** | 775.55 | 773.41 | 8 | 0.83 |
| ***Sgs7*** | 663.20 | 659.79 | 8 | 0.56 |
| ***Sgs3*** | 1004.24 | 992.28 | 14 | **0.046** |
| ***Sgs5*** | 1107.22 | 1102.39 | 8 | 0.29 |
| ***Sgs5bis*** | 929.55 | 926.06 | 6 | 0.32 |
| ***Sgs4*** | 1217.13 | 1211.13 | 8 | 0.15 |
| ***Eig71Ee*** | 2021.20 | 2116.73 | 8 | 0.54 |
| ***Sgs1*** | 2079.27 | 2076.58 | 8 | 0.72 |

**Table S2. Test for lineage-specific selection.** Likelihood values (-lnL) obtained for the one ratio (M0) and the free ratio models from PAML are shown. Df: degree of freedom used for the test corresponding to 2s-4 (s: number of species). P-values corresponding to the LRT tests between the two models are reported for each glue gene.

| | | Fst value (% in the distribution) | | | | | |
|---|---|---|---|---|---|---|---|
| gene | window(s) | EG-ZI | RAL-ZI | FR-ZI | EG-FR | EG-RAL | RAL-FR |
| *Sgs5* | 3R 13421000-13422000 | 0.16 (51.6%) | 0.12 (32.6%) | 0.13 (22.3%) | 0.035 (41.4%) | 0.06 (62.3%) | 0.039 (60.9%) |
| *Sgs5bis* | 3R 13420000-13421000 | 0.089 (20.3%) | 0.14 (42.1%) | 0.24 (63.3%) | 0.049 (50.8%) | -0.010 (7.8%) | 0.050 (70.2%) |
| *Sgs7-Sgs8* | 3L 11502000-11503000 | 0.26 (78.0%) | 0.35 (92.8%) | 0.41 (92.3%) | 0.032 (39.7%) | 0.015 (26.2%) | 0.019 (35.2%) |
| *Sgs7* | 3L 11503000-11504000 | 0.28 (81.5) | 0.30 (87.7%) | 0.41 (92.4%) | 0.014 (26.6%) | -0.02 (3.4%) | 0.04 (61.9%) |
| *Sgs3* | 3L 11505000-11506000 | 0.53 (98.4%) | 0.59 (99.4%) | 0.71 (99.7%) | 0.081 (67.2%) | -0.025 (2.3%) | 0.053 (72.2%) |
| *Sgs3* | 3L 11506000-11507000 | 0.25 (76.0%) | 0.28 (84.8%) | 0.36 (87.3%) | 0.050 (53.7%) | 0.0070 (19.3%) | 0.025 (43.5%) |
| *Sgs1* | 2L 4937000-4938000 | 0.064 (11.1%) | (0.15) 45.3% | 0.16 (35.3%) | 0.065 (59.9%) | 0.064 (65.0%) | 0.020 (37.5%) |
| *Sgs1* | 2L 4938000-4939000 | 0.044 (5.3%) | 0.14 (42.9%) | 0.24 (64.5%) | 0.17 (90.5%) | 0.082 (73.7%) | 0.014 (27.4%) |
| *Sgs1* | 2L 4939000-4940000 | 0.18 (58.6%) | 0.25 (80.6%) | 0.12 (18.2%) | 0.00 (15.2%) | 0.11 (83.8%) | 0.048 (69.3%) |
| *Sgs1* | 2L 4940000-4941000 | -0.077 (0.1%) | 0.091 (17.7%) | 0.093 (9.4%) | 0.18 (91.0%) | 0.12 (84.6%) | -0.0055 (2.3%) |
| *Sgs1* | 2L 4941000-4942000 | 0.12 (34.7%) | 0.13 (38.0%) | 0.24 (62.8%) | 0.11 (78.8%) | 0.0050 (17.6%) | 0.13 (94.3%) |
| *Eig71Ee* | 3L 15647000-15648000 | 0.14 (43.8%) | 0.11 (26.0%) | 0.21 (52.8%) | 0.10 (75.9%) | 0.095 (78.6%) | 0.073 (82.8%) |
| *Eig71Ee* | 3L 15648000-15649000 | 0.12 (33.5%) | 0.13 (36.1%) | 0.18 (42.0%) | 0.19 (92.1%) | 0.17 (92.8%) | 0.039 (60.8%) |
| *Eig71Ee* | 3L 15649000-15650000 | 0.25 (75.2%) | 0.17 (57.8%) | 0.35 (85.6%) | 0.19 (92.1%) | 0.091 (77.2%) | 0.071 (81.9%) |
| *Sgs4* | X 3143000-3144000 | 0.33 (87.6%) | 0.26 (82.2%) | 0.29 (75.5%) | 0.032 (39.9%) | 0.022 (33.1%) | 0.0026 (9.4%) |
| *Sgs4* | X 3144000-3145000 | 0.30 (84.4%) | 0.24 (78.4%) | 0.25 (66.3%) | 0.071 (63.0%) | 0.14 (89.4%) | 0.058 (75.3%) |
| *Sgs4* | X 3145000-3146000 | 0.43 (95.1%) | 0.41 (96.0%) | 0.43 (94.0%) | 0.026 (35.1%) | 0.024 (35.4%) | 0.0066 (15.1%) |

**Table S3. Fst estimates for 1kbp non-overlapping genomic windows containing the glue gene coding regions (rows) and for selected pairs of Drosophila Genome Nexus populations (columns).** The percentile of the Fst estimate relative to the total Fst distribution is indicated in parentheses. "96.0%" means that the Fst value is at the highest 4th percentile among all the Fst values calculated across the whole genome for that specific population pair. EG: Egypt, FR: France, RAL: Raleigh, ZL: Zambia.

| Species | Line | *Sgs1* | *Sgs3* | *Sgs4* | *Eig71Ee* |
|---|---|---|---|---|---|
| *D. melanogaster* | A1 | 64 [16] | 37 [5] | 21 [7] | 16 [14] |
| *D. melanogaster* | A2 | 49 [16] | 29 [5] | 21 [7] | 12 [14] |
| *D. melanogaster* | A3 | 67 [16] | 38 [5] | 17 [7] | 5 [14] |
| *D. melanogaster* | A4 | 58 [16] | 38 [5] | 22 [7] | 11 [14] |
| *D. melanogaster* | A5 | 53 [16] | 37 [5] | 21 [7] | 6 [14] |
| *D. melanogaster* | A6 | 49 [16] | 35 [5] | 22 [7] | 6 [14] |
| *D. melanogaster* | A7 | NA | 35 [5] | 30 [7] | 11 [14] |
| *D. melanogaster* | AB8 | 65 [16] | 35 [5] | 20 [7] | 12 [14] |
| *D. melanogaster* | B1 | 45 [16] | 35 [5] | 30 [7] | 12 [14] |
| *D. melanogaster* | B2 | 67 [16] | 37 [5] | 19 [7] | 6 [14] |
| *D. melanogaster* | B3 | 54 [16] | 30 [5] | 22 [7] | 5 [14] |
| *D. melanogaster* | B4 | 49 [16] | 37 [5] | 22 [7] | 11 [14] |
| *D. melanogaster* | B6 | 55 [16] | 37 [5] | 20 [7] | 5 [14] |
| *D. melanogaster* | ISO | 62 [16] | 37 [5] | 20 [7] | 11 [14] |
| *D. melanogaster* | ORE | 55 [16] | 37 [5] | 19 [7] | 11 [14] |
| *D. mauritiana* | mau12 | 36 [16] | 13 [20] | 32 [7] | ND |
| *D. sechellia* | sech25 | 34 [16] | 8 [9] | 5 [7] | 20 [12] |
| *D. simulans* | w501 | 31 [16] | ND | 7 [7] | 5 [12] |
| *D. yakuba* | NY73PB | 51 [20] | 18 [7] | 23 [7] | 6 [22] |
| *D. teissieri* | GT53w | 47 [20] | 13 [12] | 14 [7] | 6 [22] |

**Table S4. Repeat number variation in the glue genes.** The number of repeated motifs (shown) is estimated by dividing the length of the repeated region by the length of the repeated motif (in brackets). ND: the number of repeats could not be determined because the repeated motif could not be determined.

| gene | pop | $TajD_{obs}$ | 95% confidence interval | Average D | $P(D<=D_{obs})$ |
|---|---|---|---|---|---|
| ***Sgs8*** | DSPR | -0.824 | [-1.91,1.85] | -0.074 | 0.22 |
| ***Sgs8*** | ZI | -0.314 | [-1.60,2.03] | -0.083 | 0.44 |
| ***Sgs7*** | DSPR | -0.399 | [-1.16,1.50] | -0.046 | 0.53 |
| ***Sgs7*** | ZI | **-1.650** | **[-1.61,2.06]** | **-0.117** | **0.02** |
| ***Sgs3*** | DSPR | -1.322 | [-1.76,1.75] | -0.092 | 0.08 |
| ***Sgs3***[1] | DSPR | -0.948 | [-1.69,2.00] | -0.077 | 0.23 |
| ***Sgs3***[1] | ZI | -1.393 | [-1.62,1.96] | -0.072 | 0.07 |
| ***Sgs5*** | DSPR | **2.102** | **[-1.69,1.78]** | **-0.081** | **0.99** |
| ***Sgs5*** | ZI | -0.859 | [-1.57,1.92] | -0.082 | 0.21 |
| ***Sgs5bis*** | DSPR | 1.146 | [-1.85,1.82] | -0.085 | 0.90 |
| ***Sgs5bis*** | ZI | **-1.586** | **[-1.58,1.95]** | **-0.102** | **0.02** |

| | | | | | |
|---|---|---|---|---|---|
| ***Sgs4*** | DSPR | 0.099 | [-1.73,1.71] | -0.085 | 0.50 |
| ***Sgs4***[1] | DSPR | 0.860 | [-1.69,1.78] | -0.076 | 0.84 |
| ***Sgs4***[1] | ZI | -1.397 | [-1.64,1.83] | -0.090 | 0.06 |
| ***Eig71Ee*** | DSPR | -0.499 | [-1.77,1.66] | -0.094 | 0.34 |
| ***Eig71Ee***[1] | DSPR | -0.863 | [-1.72,1.70] | -0.087 | 0.20 |
| ***Eig71Ee***[1] | ZI | -0.654 | [-1.61,1.83] | -0.117 | 0.30 |
| ***Sgs1*** | DSPR | -1.419 | [-1.76,1.62] | -0.110 | 0.07 |
| ***Sgs1***[1] | DSPR | -0.943 | [-1.69,1.85] | -0.073 | 0.26 |
| ***Sgs1***[1] | ZI | **-1.507** | **[-1.60,1.86]** | **-0.105** | **0.03** |

**Table S5. Coalescent simulations for Tajima's D test significance.** $TajD_{obs}$: TajD value observed. 95% confidence interval in which TajD value is expected under the standard neutral model. Average D obtained by simulations. P(D<=$D_{obs}$): probability that $TajD_{obs}$ is higher than expected under the standard neutral model.

### Datasets and scripts

(available on Dryad, doi:10.5061/dryad.v6wwpzgwk, temporary link for the review process: https://datadryad.org/stash/share/KN532B9hnGG_v17jpC2ZqyM4UQSnxK8GS1BkaZgp3oc)

**Dataset1.ods. Raw data**
Sheet1: Stock_information: Information of the fly lines used for the analysis.
Sheet2: adhesion_test: Results of the adhesion assay performed on pupae used in the analyses (including adhesion tests that were excluded from further analysis).
Sheet3: glue_area: Pupa-substrate contact area measurements.
Sheet4: codon_usage_and_expression: Expression level and codon adaptation index for genes expressed in salivary glands of wandering larva.
Sheet5: PAML_dN_values: synonymous and non synonymous substitution rates obtained with PAML under M0 and free ratio models.
Sheet6: FEL_results: raw results obtained by the Fixed Effect Likelihood method implemented in HyPhy via the datamonkey web platform.

**fasta_alignments.zip:** Sequence alignments used for the different analyses.

**scripts.zip:** folder containing scripts used to do the analyses.

**circos_raw_data.zip:** data used to produce circos plot figures.

**glue_prints.zip:** folder containing pictures of prints left by pupae after detachment.
